# Effect of initial body orientation on escape probability in prey fish escaping from predators

**DOI:** 10.1101/134494

**Authors:** Hibiki Kimura, Yuuki Kawabata

**Affiliations:** Graduate School of Fisheries and Environmental Sciences, Nagasaki University, Bunkyo-machi, Nagasaki 852-8521, Japan

**Keywords:** Attack angle, C-start, Escape response, Fast-start, Kinematics, Predator-prey interaction

## Abstract

Since the escape response is crucial to survival and hence to the fitness of species, several studies have attempted to elucidate the kinematic and behavioral components of the response that affect evasion outcome. The prey’s body orientation relative to a predator at the onset of the escape response (initial orientation) could affect evasion outcome, because the turn angle and its duration before the initiation of escape locomotion would be smaller when the initial orientation is more away from the predator. We tested this hypothesis by recording the escape responses of juvenile red sea bream (*Pagrus major*) in response to the predatory scorpion fish (*Sebastiscus marmoratus*) using a high-speed video camera. Our results show that an increased initial orientation (i.e., more away from the predator) increases escape probability. Our results also indicate that an increase in the initial orientation decreases the turn angle and its duration. The flight initiation distance tends to be small when the initial orientation is away from the predator, suggesting that the prey might have a blind zone of sensory perception. These findings highlight the importance of incorporating initial orientation into both empirical and theoretical studies of the kinematics of predator-prey interactions.

**Summary statement:** Our predator-prey experiments reveal that the prey’s initial body orientation relative to a predator affects the prey’s turn angle and its duration, and consequently affects escape probability.

## Introduction

When exposed to sudden predation threats, most animals exhibit escape responses that include turning swiftly and accelerating forward (Bulbert et al., 2015; Camhi et al., 1978; Webb, 1986). Since the escape response is crucial to survival and hence to the fitness of the species, numerous studies have been conducted to elucidate the environmental and internal factors that affect the behavioral and kinematic components of the escape response (e.g., flight initiation distance, escape trajectory, turning speed, acceleration, etc.) (Bateman and Fleming, 2014; Cooper, 2006; Cooper et al., 2007; Domenici, 2010; Meager et al., 2006). Most of these studies, however, have used artificial stimuli to elicit the escape response, and thus knowledge of the importance of different components of the response in the context of avoiding real predators is still limited.

Previous theoretical studies have shown that the outcome of the escape response is dependent on the flight initiation distance, predator and prey speeds, and the escape trajectory (Arnott et al., 1999; Broom and Ruxton, 2005; Domenici, 2002; Weihs and Webb, 1984). Interestingly, however, these studies have not incorporated the prey’s initial body orientation with respect to the predator (hereafter, initial orientation) and the prey’s turning speed, despite the fact that turning requires additional time prior to the initiation of escape locomotion (King and Comer, 1996), and that initial orientation affects the turn angle (Cooper and Sherbrooke, 2016; Eaton and Emberley, 1991; Kawabata et al., 2016). Empirical studies show that turning speed, as well as the above variables, affects predator evasion (Dangles et al., 2006; Scharf et al., 2003; Stewart et al., 2013; Walker et al., 2005); however, as far as we aware, except for one study (Stewart et al., 2013), no research has been conducted on the effect of initial orientation on escape probability.

The C-start escape response of fish and amphibian larvae is one of the most well-studied escape responses in animals (Domenici and Blake, 1997; Eaton et al., 2001). The C-start escape response is composed of three distinct stages based on kinematics: the initial bend (stage 1), the return tail flip (stage 2), and then continuous swimming or coasting (stage 3) (Domenici and Blake, 1997; Weihs, 1973). It has been shown that flight initiation distance, escape speed, turning speed, and escape trajectory affect evasion outcome (Scharf et al., 2003; Stewart et al., 2013; Walker et al., 2005), but the effect of initial orientation remains to be elucidated. The objectives of our study were to determine whether initial orientation affects evasion outcome, and if so, to investigate the mechanisms involved. To achieve these objectives, we recorded the escape responses of juvenile red sea bream [*Pagrus major* (Temminck & Schlegel, 1843)] in response to the predatory scorpion fish [*Sebastiscus marmoratus* (Cuvier, 1829)] using a high-speed video camera. Since the fish could have spatial bias in detecting the attacking predator (e.g., a sensory blind zone), the effect of initial orientation on the response parameters was also examined. The specific questions addressed were as follows: (1) does an increase in the initial orientation of prey fish (more opposite from the direction of the predator) increase escape probability?; (2) does an increase in the initial orientation decrease the turn angle and its duration?; and (3) does the initial orientation affect responsiveness and flight initiation distance?

## Results

In general, the predator [*S. marmoratus*, 149.9±17.0 (mean±s.d.) mm total length (TL), *n*=7] approached the prey (*P. major*, 56.1±9.6 mm TL, *n*=46) and then attacked it by opening its mouth. The kinematic stages in which the prey were captured are summarized in Fig. 1. The most prey individuals (43/46: 93%) showed escape responses (C-start), but three (3/46: 7%) did not show responses and were captured by predators. Of the 43 prey that showed escape responses, 19 (19/43: 44%) were captured by predators during stage 1. Of the 24 prey that survived until the end of the stage 1, four (4/24: 17%) were captured by the end of stage 2. No fish were captured during stage 3. Of the total number of prey captured (26), 22 (22/26: 85%) were captured by the end of stage 1. These results indicate that stage 1 is the most critical period for *P. major* to escape from the attack of *S. marmoratus*.

**Fig. 1.**
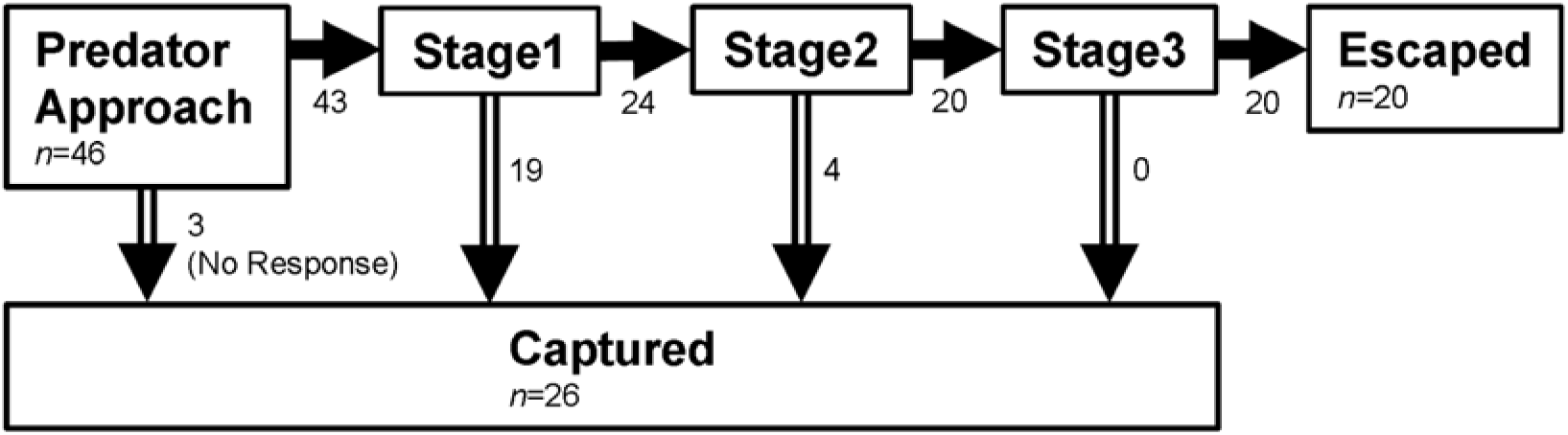
Diagram showing the kinematic stages in which the prey were captured.

The frequency distribution of the initial orientation and the initial orientation–escape probability relationship are shown in Fig. 2. The frequency of initial orientation at 120-180° was lower than at 0-120° (Fig. 2A). Escape probability was highest in the 120-150° initial orientation bin, although 95% confidence intervals based on binomial distributions suggest that there were no significant differences among the different initial orientation bins (Fig. 2B).

**Fig. 2.**
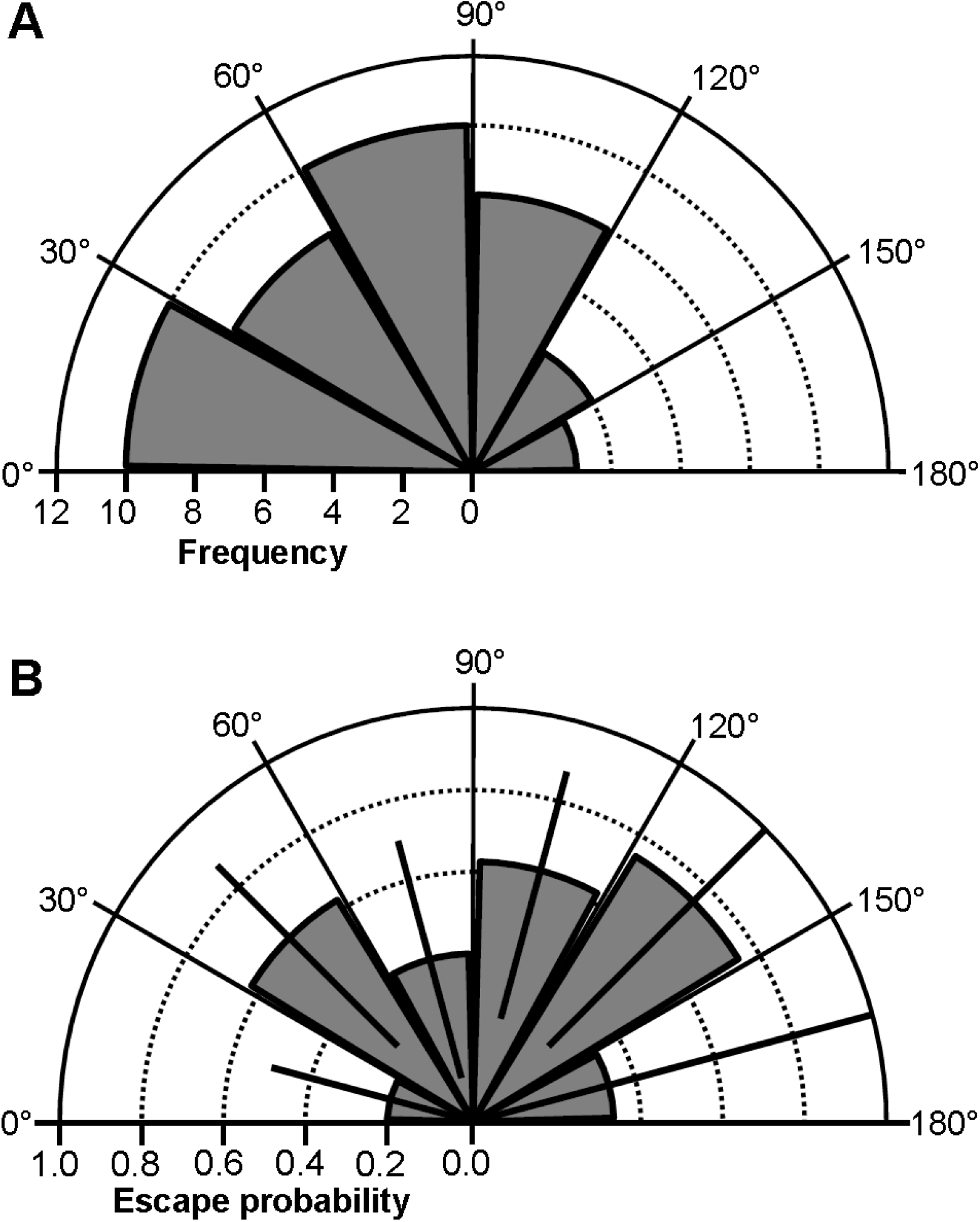
(A) Frequency distribution of initial orientations. (B) Relationship between initial orientation and escape probability. The error bars represent 95% confidence intervals, estimated by assuming binomial distributions.

Differences in the parameters (initial orientation, flight initiation distance, and predator speed) between the successful (escaped) and unsuccessful (captured) escapes are shown in Table 1. The initial orientation of the successful escapes (79.7±43.5°) was larger than that of the unsuccessful ones (64.2±51.0°), and the larger initial orientation significantly increased escape probability (Fig. 3; LR-test, χ^2^=5.30, d.f.=1, *P*<0.05). The odds ratio indicates that a 48.0° (1 s.d.) increase in initial orientation increased the escape probability 2.52 times. Increases in flight initiation distance also significantly increased escape probability (Fig. 3; LR test, χ^2^=17.98, d.f.=1, *P*<0.01), but the effect of predator speed was insignificant (LR test, χ^2^=0.23, d.f.=1, *P*=0.63). The odds ratio of flight initiation distance indicates that an increase of 31.6 mm (1 s.d.) increased the escape probability 6.47 times.

**Table 1.**
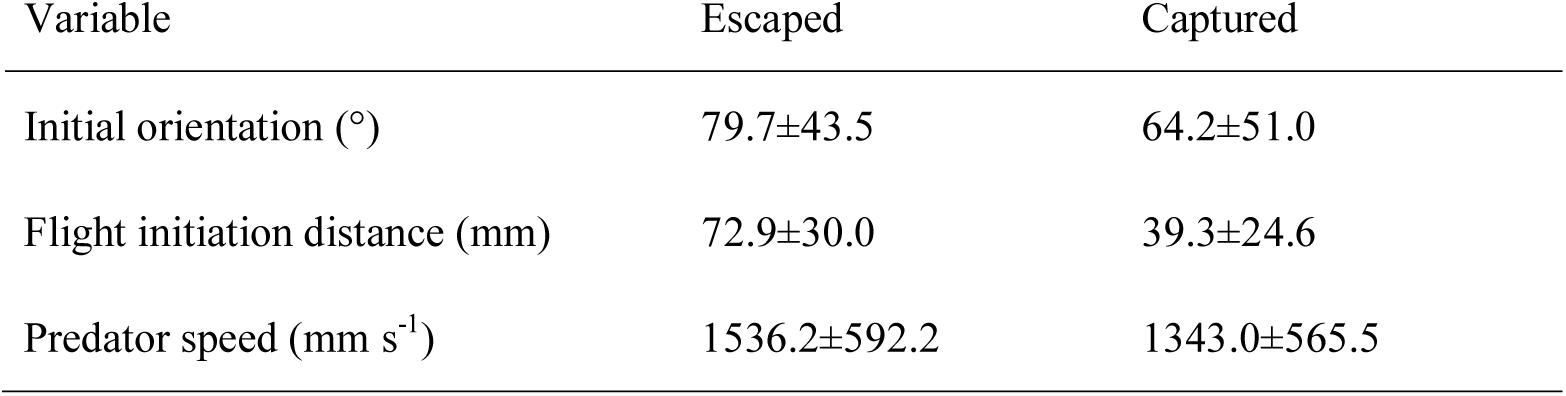
Comparisons of variables between successful (escaped) and unsuccessful (captured) escapes.

| Variable | Escaped | Captured |
| --- | --- | --- |
| Initial orientation (°) | 79.7±43.5 | 64.2±51.0 |
| Flight initiation distance (mm) | 72.9±30.0 | 39.3±24.6 |
| Predator speed (mm s<sup>-1</sup>) | 1536.2±592.2 | 1343.0±565.5 |

**Fig. 3.**
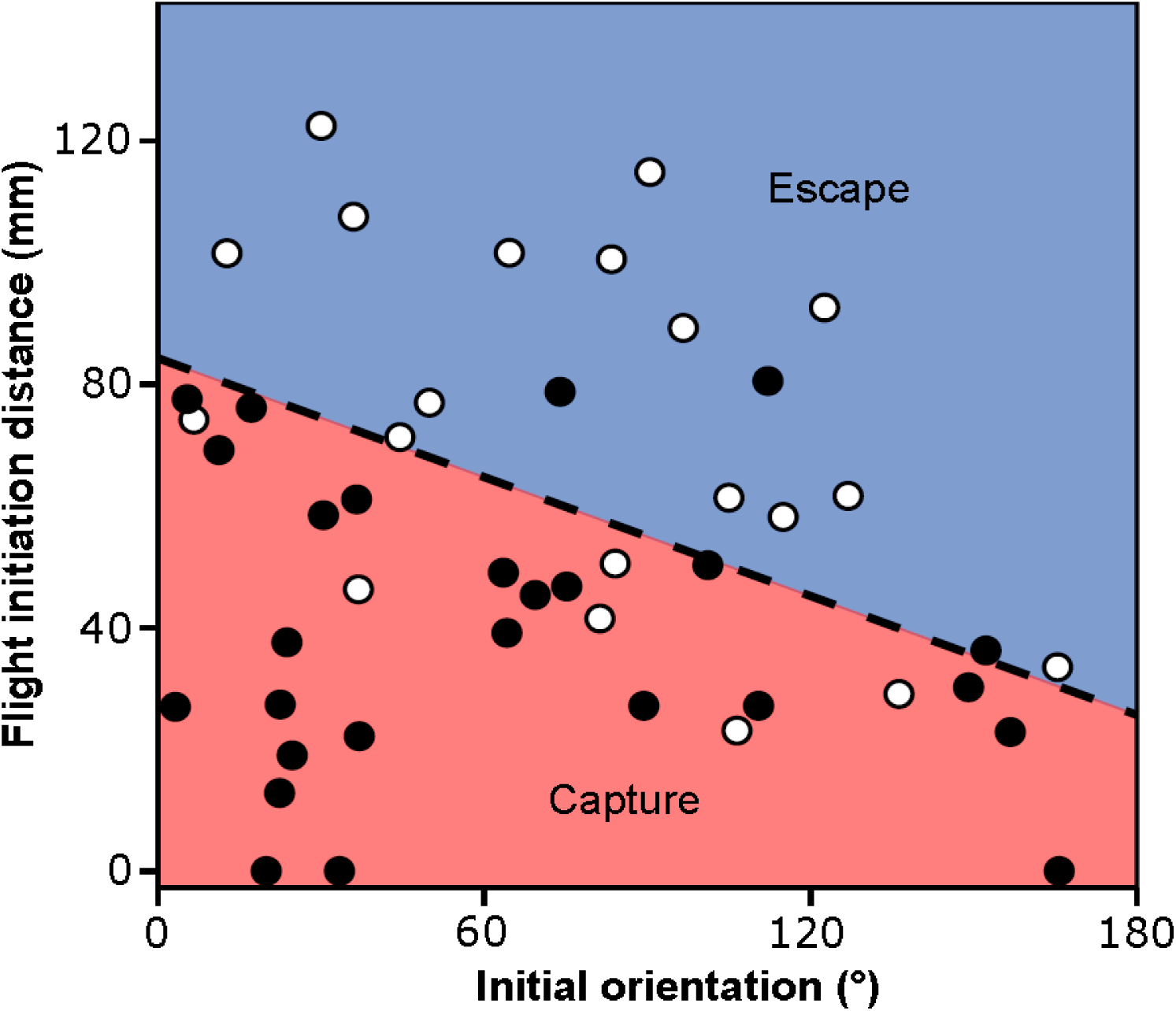
Effect of initial orientation and flight initiation distance on survival probability. Open circles are indicative of successful escape from predator’s attack and filled circles are indicative of captured by predator’s attack. The dashed line represents the 50% escape probability estimated from the generalized linear mixed model (GLMM). All the prey fish were used in this analysis (*n*=46).

There were negative relationships between initial orientation and turn angle (Fig. 4A; *R*=−0.61, *n=*24, *P*<0.01), and between the initial orientation and turn duration (Fig. 4B; *R*=−0.41, *n*=24, *P*<0.05); the effect of the initial orientation on turn angle and turn duration was significant (turn angle: LMM, *F*_1,18.5_=22.88, *P*<0.01; turn duration: LMM, *F*_1,16.9_=29.56, *P*<0.01). Additionally, there was a significant positive relationship between the turn angle and its duration (*R*=0.53, *n*=24, *P*<0.01). These results indicate that the turn angle and its duration were larger when the initial orientation was more toward the predator, and smaller when the initial orientation was more away from the predator.

**Fig. 4.**
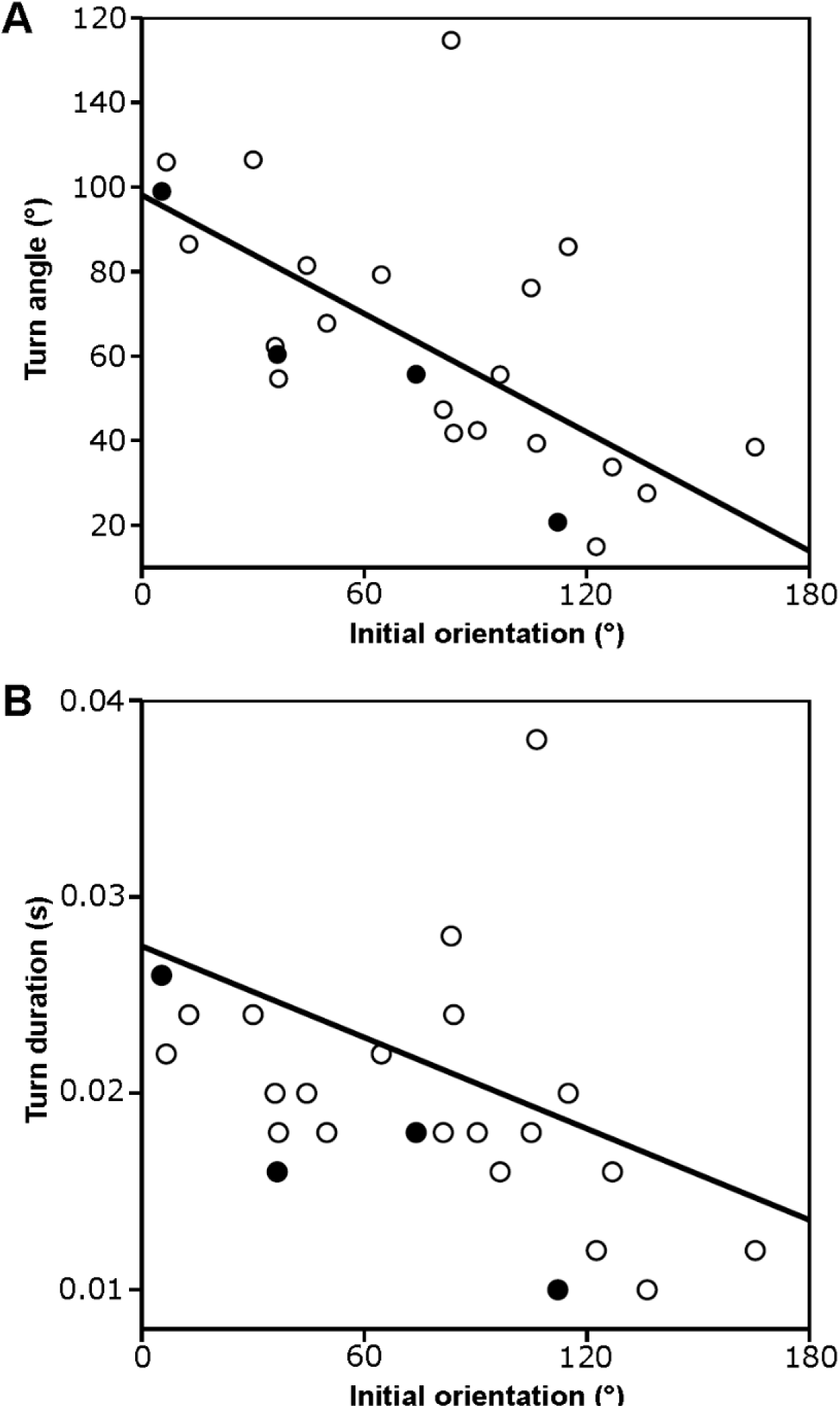
(A) Relationship between initial orientation and turn angle. (B) Relationship between initial orientation and turn duration. Open circles are indicative of successful escape from predator’s attack and filled circles are indicative of captured by predator’s attack. Prey fish that survived until the end of stage 1 were used in this analysis (*n*=24).

There was no observable pattern in the initial orientations of the three prey individuals that did not show escape responses (19.9, 33.4, and 165.7°). The flight initiation distance tended to be shorter when the initial orientation was away from predators (about 150-180°; Fig. 5), although this tendency (the effect of initial orientation on flight initiation distance) was not statistically significant (GAMM, *F*=2.10, estimated d.f.=2.35, estimated residual d.f.=40.92, *P*=0.09). Predator speed significantly increased the flight initiation distance of the prey (GAMM, *F*=4.82, estimated d.f.=1.73, estimated residual d.f.=40.92, *P*<0.05).

**Fig. 5.**
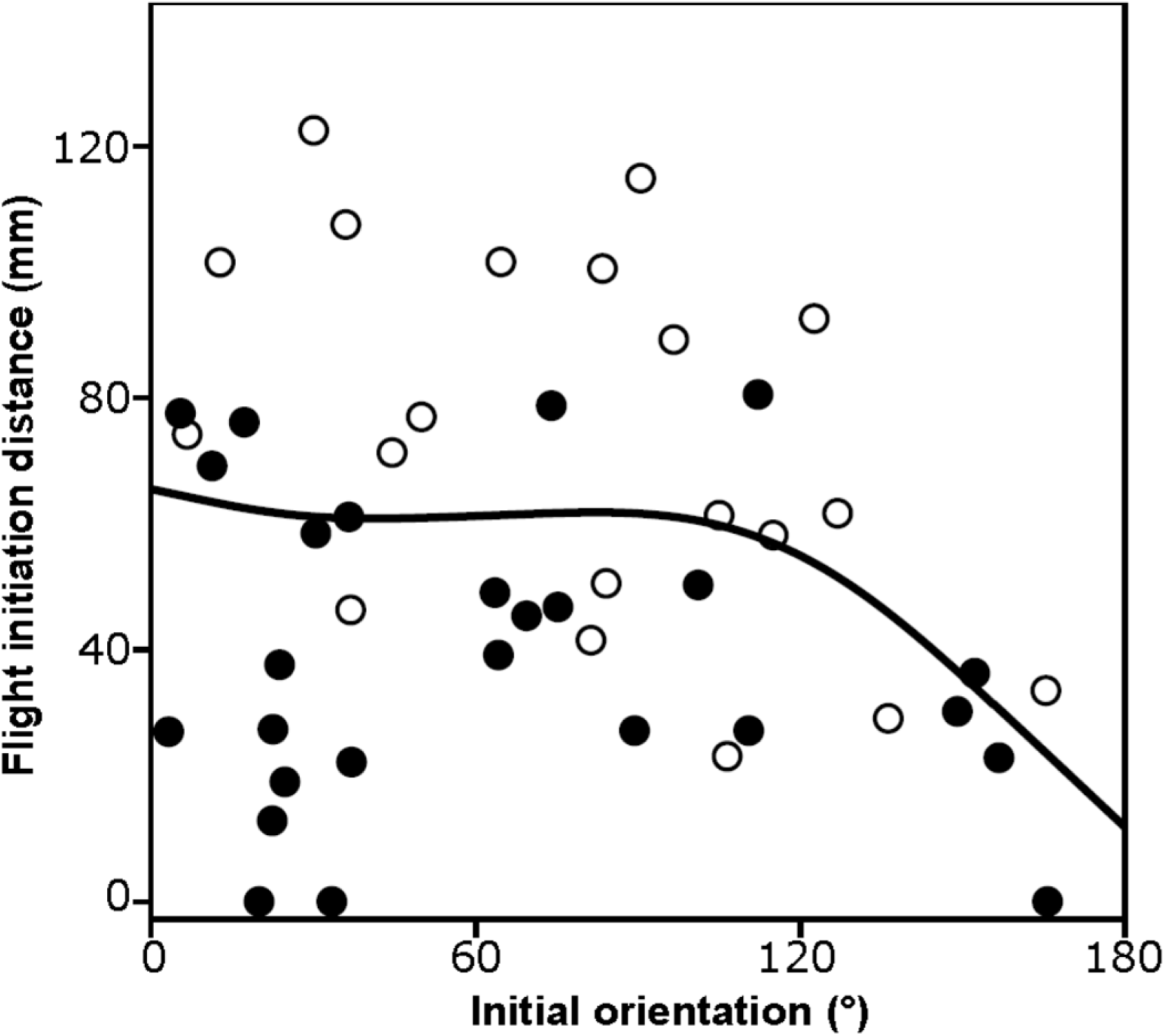
Effect of initial orientation on flight initiation distance. The line was estimated by the generalized additive mixed model (GAMM), in which the predator speed was regarded as its mean value (1427 m s^−1^). All the prey fish were used in this analysis (*n*=46).

## Discussion

Our results clearly show that an increase in the initial orientation (i.e., more fully away from the predator) increases the probability that *P. major* juveniles will escape from the predatory strikes of *S. marmoratus* (Fig. 3). This result is inconsistent with a study of zebrafish larvae evading adult zebrafish (Stewart et al., 2013), in which escape probabilities were not significantly different among six different initial orientation bins. This discrepancy could be attributed to the different statistical methods used in the two studies, or to species-specific/ontogenetic differences. In the study of the zebrafish larvae, the initial orientation values were categorized into six bins and escape probability was compared among the different bins by calculating 95% confidence intervals. In that analysis, the effects of other variables (i.e., flight initiation distance and predator speed) were not offset. By contrast, our study used a generalized linear mixed model (GLMM) with no binning of the initial orientation values, and it included the other possible variables in the model to offset their variation. In fact, when we analyzed the effect of initial orientation in the same manner as in the study of the zebrafish larvae, the effect of initial orientation on escape probability became statistically insignificant (Fig. 2B). In light of these facts, the binning procedures and/or the large variation in the other variables may have masked the actual effect of initial orientation, and thus initial orientation could actually be a crucial parameter for predator evasion in other fishes as well.

Our results also show that an increase in the initial orientation decreases the turn angle and its duration (Fig. 4). This initial orientation–turn angle relationship is consistent with studies of many animal taxa (e.g., other fish, frogs, cockroaches, and lizards) (Camhi and Tom, 1978; Cooper and Sherbrooke, 2016; Eaton and Emberley, 1991; King and Comer, 1996). Although a limited number of studies have examined the relationship between turn angle and its duration or between initial orientation and turn duration, it is natural to assume that a larger turn angle requires a longer duration, as has been shown in this study and in a study on frogs (King and Comer, 1996). C-starts and other escape responses start from initial turns, followed by escape locomotion; during the initial turns, the animals do not move large distances but stay close to their initial positions (Camhi et al., 1978; Domenici and Blake, 1997; King and Comer, 1996; Tauber and Camhi, 1995). Therefore, predators would be able to approach prey animals during these initial turns. It is thus likely that initial orientation-mediated turn angle changes affect escape probability by changing the time available for the predator to approach the prey before the initiation of escape locomotion.

The flight initiation distance tended to be smaller when the predator approached the prey from behind (Fig. 5). This might be related to a sensory blind zone in the prey. The C-start escape response is triggered by either visual (Dill, 1974), mechanical (Umeda et al., 2016), or sound stimuli (Domenici and Batty, 1997). When it is triggered by visual stimuli there would be a blind zone for the prey (Domenici, 2002; Tyrrell and Fernandez-Juricic, 2015). On the other hand, the lateral line (mechanosensory system) is distributed throughout the body (Dijkgraaf, 1963; Kasumyan, 2003), which may allow 360° perception without any spatial bias. It is also unlikely that there is a spatial bias in detecting sound stimuli. It is thus possible that the *P. major* juveniles relied mainly on visual senses to perform escape responses, and thus the flight initiation distance tended to be smaller when the initial orientation was away from the predator. Further research is clearly needed to clarify the relationship between initial orientation and flight initiation distance, as well as the underlying sensory mechanisms.

Considering the time for turning and the sensory blind zone, the optimal initial orientation for prey to escape from predators might be near the edge of the maximum perception range, since this would require a relatively shorter time for turning before escape locomotion, and would allow the prey to respond to the predator’s strike from a great enough distance. This hypothesis is consistent with the initial orientation–escape probability relationship, in which the maximum escape probability occurred around 120-150° (Fig. 2B). However, the frequency of the initial orientation was not highest around this range: the frequency at 120-180° was smaller than that at 0-120° (Fig. 2A). Because we used naïve hatchery-reared fish that had not experienced any predators, the prey might not have recognized the predator as dangerous, and thus the prey did not adjust the initial orientation in advance. It has been shown that black goby change their posture when a weak stimulus is presented before the strong stimulation that finally elicits an escape response (Turesson et al., 2009). Therefore, prey animals that recognize a predator in advance may adjust their initial orientation to maximize their escape probability, although we should note that predators may also adjust the attack angle (i.e., initial orientation) to maximize predation probability (Webb and Skadsen, 1980).

Different geometrical models have been proposed to explain the factors affecting escape probability and/or the escape trajectory (Arnott et al., 1999; Corcoran and Conner, 2016; Domenici, 2002; Howland, 1974; Weihs and Webb, 1984), but none of these models have incorporated initial orientation. Furthermore, initial orientation has not been considered in many empirical studies of predator-prey interactions (e.g., Dangles et al., 2006; Fuiman, 1993; Scharf et al., 2003; Walker et al., 2005). Our results clearly show that initial orientation affects escape probability, and that it can affect flight initiation distance. These findings highlight the importance of incorporating data on initial orientation into both theoretical and empirical studies of predator-prey interactions.

## Materials and Methods

### Ethics statement

Animal care and experimental procedures were approved by the Animal Care and Use Committee of the Institute for East China Sea Research, Nagasaki University (Permit no. ECSER15-12), in accordance with the Regulations of the Animal Care and Use Committee of Nagasaki University.

### Fish samples

Hatchery-reared *P. major* (*n*=151) were utilized as prey fish in this study. All individual *P. major* were provided from commercial hatcheries, and were kept in three 200 L polycarbonate tanks at the Institute for East China Sea Research, Nagasaki University, Japan. They were fed with commercial pellets (Otohime C2, Marubeni Nisshin Feed Co., Ltd., Tokyo, Japan) twice a day.

As predators, we used *S. marmoratus* (*n*=7), which is a common reef predator around the coast of Japan. *S. marmoratus* usually employs a “stalk-and-attack” tactic. All *S. marmoratus* were collected by hook-and-line around Nagasaki prefecture, Japan. The collected *S. marmoratus* were kept in a glass aquarium (1200×450×450 mm) before the start of the experiment. They were standardly fed krill once every 2-4 days.

The position of the center of mass (CM) for *P. major* was estimated by hanging dead fish (54.3±3.3 mm TL, *n*=10) from two different points using a suture and needle (Lefrancois et al., 2005). The CM position from the tip of the head was estimated as 0.34±0.01 TL.

### Experimental procedure

Experiments were performed in a glass aquarium (900×600×300 mm) with seawater to a depth of 100 mm. The water temperature during the experiments was 23.1±0.9°C. White plastic plates with grid lines were placed on the bottom and three sides of the tank; one side (900×300 mm) of the tank was left transparent to record the side view of the fish. A preliminary experiment showed that *S. marmoratus* actively fed in low light conditions, so two LED bulbs covered with red cellophane were used to illuminate the tank. The light intensity was maintained at 54 lux. Two synchronized high-speed video cameras (HAS-L1, Ditect Co., Tokyo, Japan) were used to record dorsal and side views of the fish simultaneously. (Note that we only used the dorsal views in this study.)

An individual *S. marmoratus* starved for at least 24 h was first introduced into the experimental tank and allowed to acclimate for 30 min. An individual *P. major* was then introduced into a PVC pipe (60 mm diameter) with 112 small holes (3 mm diameter) set in the center of the tank, and acclimated for 15 min. The 15-min period was chosen because a preliminary experiment showed that the fish settled down and opercular beat frequency recovered to the basal level within at most 15 min. After the acclimation period, the trial was started by slowly removing the PVC pipe to release the *P. major*. When *S. marmoratus* attacked the *P. major*, we recorded the movements of both predator and prey using the high-speed video cameras. If *S. marmoratus* did not show any predatory movements for 20 min, the trial was ended. Seven *S. marmoratus* were repeatedly used, but each *P. major* was used only once.

### Analysis of video sequences

Because the vertical displacements of both fishes were negligible, we only used the dorsal video views in our analyses. Before measuring the kinematic and behavioral variables, we noted the kinematic stage in which each prey was captured. The escape response of *P. major* and the predatory strike of *S. marmoratus* were then analyzed frame by frame using Dipp-Motion Pro 2D (Ditect Co., Tokyo, Japan). The CM and the tip of the snout of *P. major*, and the tip of the snout of *S. marmoratus*, were digitized in each frame, and the following variables were calculated:

Flight initiation distance: the distance between the predator’s snout and the prey’s CM at the onset of stage 1 (Fig. 6, D0). Initial orientation (°): the angle between the line passing through the predator’s snout and the prey’s CM, and the line passing through the prey’s CM and the prey’s snout at the onset of the stage 1 (Fig. 6, A0). Turn angle (°): the angle between the line passing through the prey’s CM and the prey’s snout at the onset of stage 1, and the line passing through the prey’s CM and the prey’s snout at the onset of the return tail flip (Fig. 6, A1). Turn duration (s): the time between the onset of stage 1 and the onset of the return tail flip. Predator speed (mm s^-1^): the cumulative distance the predator’s snout moves during the period between the onset of stage 1 and 0.01 s before the onset of stage 1, multiplied by 100.

**Fig. 6.**
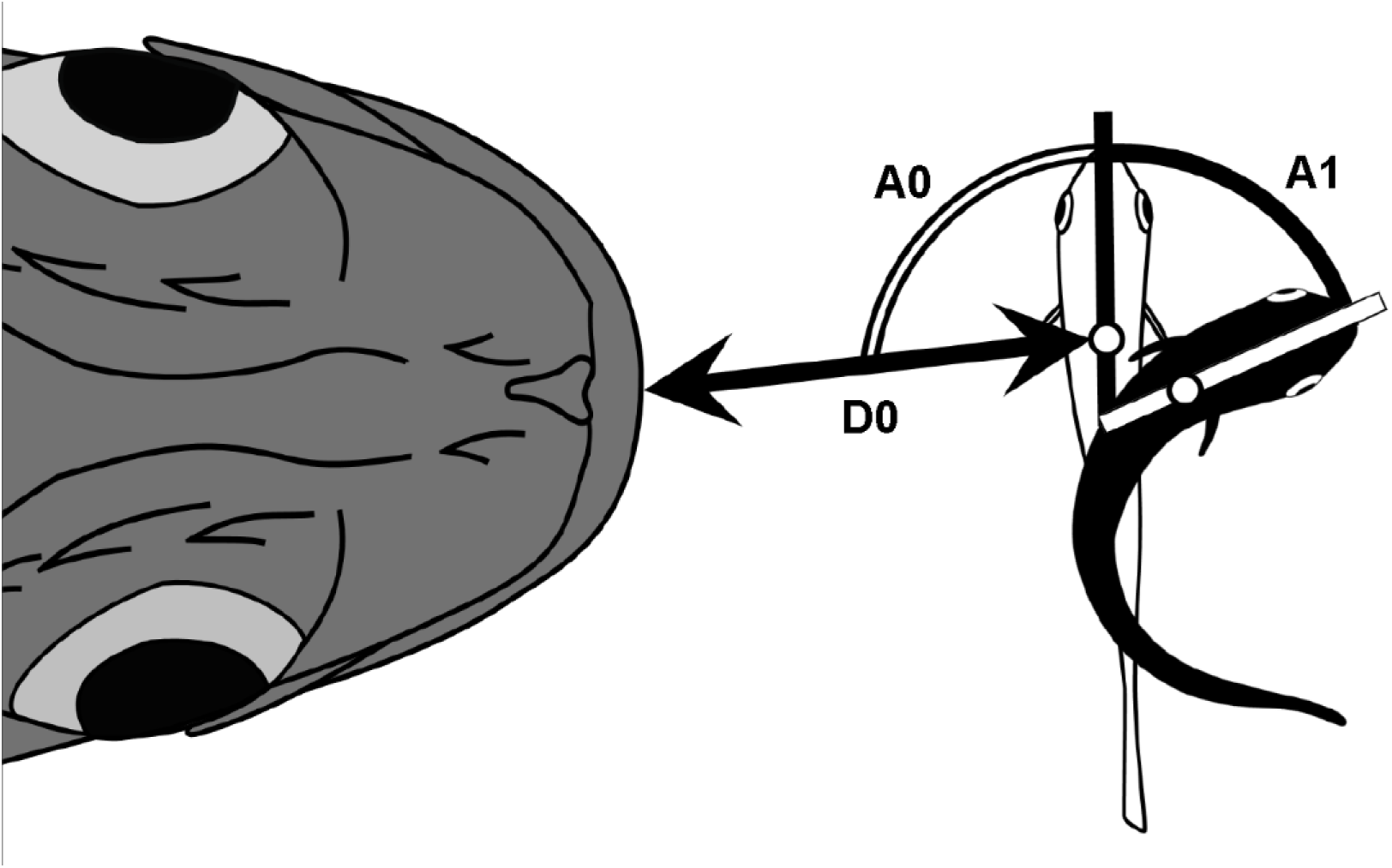
Schematic drawing of measured variables. The position of the prey at the onset of the escape response is shown as an unfilled fish, and the position at the end of stage 1 is shown as a filled fish. Unfilled circles represent the prey’s center of mass (CM). D0, flight initiation distance; A0, initial orientation; A1, turn angle.

When prey fish did not show escape responses (*n*=3, Fig. 1), the flight initiation distance was regarded as 0. The initial orientation relative to a predator was calculated at the onset of the predator’s strike. The predator speed was calculated during the period between the time of capture and 0.01 s before the time of capture.

### Statistical analyses

Of the 151 digital films recorded, 46 were used for the data analyses. First, fish that were not sufficiently far from the wall (more than one total length) were omitted from the analysis to eliminate possible wall effects (Eaton and Emberley, 1991). Second, only fish that initiated an escape response from a state of rest were used in the analysis (we excluded cases where *S. marmoratus* chased *P. major* that were already swimming).

To test the hypothesis that initial orientation affects escape probability, the effects of initial orientation on escape probability were evaluated using a generalized linear mixed model (GLMM) with a binomial error distribution and a logit link function (Zuur et al., 2009). All the fish were used in this analysis (*n*=46). Success and failure of predator evasion were designated as 1 and 0, respectively, and used as the objective variable. Initial orientation, flight initiation distance, and predator speed were considered as explanatory variables; flight initiation distance and predator speed were included in the model because these variables significantly affected escape probability in previous studies (Dangles et al., 2006; Stewart et al., 2013; Walker et al., 2005). Predator ID was also included as a random factor because unknown predator abilities may affect the evasion outcome. The significance of the explanatory variables was then assessed by progressively removing them from the model and comparing the change in deviance using the likelihood ratio test with a χ^2^ distribution (LR test). The final model for estimating the escape probability was also determined by progressively removing the explanatory variables when the variables were not significant in the LR test.

The second hypothesis, that initial orientation decreases the turn angle and its duration, was evaluated using a linear mixed model (LMM) (Grafen and Hails, 2002; Zuur et al., 2009). Prey fish that survived until the end of stage 1 were used in this analysis (*n*=24, Fig. 1). Turn angle or turn duration was used as the objective variable, and initial orientation was considered as the explanatory variable. Predator ID was also included as a random factor. The significance of the explanatory variable was assessed by the *F* test.

Prey animals can have spatial bias in detecting an attacking predator (e.g., from a sensory blind zone) (Domenici, 2002; Tyrrell and Fernandez-Juricic, 2015). Therefore, we examined whether the initial orientation affected the responsive parameters. Because a majority of the prey (43/46, 93%) showed escape responses, we could not conduct any statistical analysis regarding responsiveness. Instead, we examined whether initial orientation affected the flight initiation distance using a generalized additive mixed model (GAMM) with a normal error distribution and an identity link function (Zuur et al., 2009). The GAMM was used because flight initiation distance is likely to change in response to changes in initial orientation in a non-linear fashion due to the sensory blind zone. All the fish were used in this analysis (*n*=46). Flight initiation distance was used as the objective variable, and initial orientation and predator speed were considered as explanatory variables. Predator ID was also included as a random factor. The significance of the explanatory variables was assessed by the *F* test. All the analyses were carried out using R 3.3.2 (The R Foundation for Statistical Computing, Vienna, Austria) with the package *lme4* for GLMM and LMM, and the package *gamm4* for GAMM.

## Acknowledgements

We thank N. Nishiumi, I. Nakamura and A. Matsuo for their help with the experiment, G. N. Nishihara for his advice on the statistical analyses, and K. Tsurui and Y. Iwatani for kindly allowing us to use their high-speed video camera. We also thank Nagasaki Takashima Fisheries Center for kindly providing hatchery-reared red seabream.

### Competing interests

No competing interests to declare.

### Author contributions

Y.K. designed the experiment. H.K. collected the data. H.K. and Y.K. analyzed the data.

Y.K. wrote the manuscript.

### Funding

This research received no specific grant from any funding agency in the public, commercial, or not-for-profit sectors.

### Data availability

Measured variables (initial orientation, flight initiation distance, turn angle, turn duration, and predator speed), predator ID, and evasion outcome from 46 predator-prey interactions are available as supplementary information in Table S1.

